# Sex differences in corticostriatal regulation of alcohol drinking following chronic intermittent alcohol exposure

**DOI:** 10.64898/2026.11.24.769022

**Authors:** Kelly A. Hewitt, Skylar E. Nicholson, Danielle Logan, Emily Qualls-Creekmore, Angela M. Henricks

**Affiliations:** Department of Psychology, Washington State University, Pullman, WA 99164, USA; Integrative Physiology and Neuroscience, Washington State University, Pullman, WA 99164, USA

**Keywords:** Alcohol withdrawal, chemogenetics, corticostriatal circuit, elevated plus maze, sex-differences

## Abstract

**Background:** Sex differences in alcohol use disorder (AUD) suggest that the neural circuits regulating alcohol intake may differ between males and females. We examined whether infralimbic neurons projecting to the nucleus accumbens shell (IL→NAcSh) regulate alcohol self-administration and anxiety-related behavior following chronic intermittent alcohol (CIA) exposure.

**Methods:** Male and female Sprague–Dawley rats were trained to self-administer 10% alcohol before receiving excitatory, inhibitory or control designer receptors exclusively activated by designer drugs (DREADDs) in NAcSh–projecting IL neurons. Rats then underwent six weeks of CIA vapor or room air exposure. Alcohol self-administration and elevated plus maze behavior was then assessed after deschloroclozapine pretreatment.

**Results:** During acute withdrawal, IL→NAcSh inhibition increased alcohol intake in CIA-exposed males, while excitation increased intake in room air control males. Neither manipulation detectably altered alcohol intake in females. In the elevated plus maze, excitation reduced open-arm time and increased closed-arm time in males, while both excitation and inhibition reduced open-arm time in females.

**Conclusions:** Activity in NAcSh–projecting IL neurons appears to constrain alcohol intake during acute withdrawal in CIA-exposed males. Increasing activity in these neurons did not reduce intake in this group. The divergent effects on EPM behavior and drinking suggest that this pathway’s contributions to anxiety-like behavior and alcohol self-administration are partly dissociable. The absence of detectable effects on drinking in females highlights the need to identify additional pathways that regulate dependence-related alcohol use in females. Better defining the circuits that regulate withdrawal-induced drinking in males and females may guide the development of more effective, sex-informed circuit-based therapies for AUD.

## INTRODUCTION

While men consume more alcohol and are more likely to be diagnosed with alcohol use disorder (AUD) compared with women, alcohol use among women has increased substantially over the past several years (Erol & Karpyak, 2015; Grant et al., 2015; White, 2020). Further, women often differ from men in the factors that motivate alcohol use, the progression of dependence, and their response to currently available pharmacotherapies (Canidate et al., 2017; Garbutt et al., 2005). Historically, however, the vast majority of preclinical neuroscience studies investigating AUD were conducted almost exclusively in male animals, limiting our understanding of whether alcohol misuse is driven by the same neurobiological mechanisms in males and females (Karp & Reavey, 2019). Identifying both shared and sex-specific mechanisms underlying AUD is essential for developing more effective and broadly applicable treatment approaches.

Projections from the medial prefrontal cortex (mPFC) to nucleus accumbens (NAc) have emerged as a particularly important target for regulating alcohol-related behaviors. The mPFC provides executive control over reward-seeking by integrating contextual information, expected outcomes and behavioral goals, whereas the NAc integrates these cortical inputs with limbic and dopaminergic inputs to guide motivated behavior (Haber, 2016; Marchant et al., 2015). Chronic alcohol exposure disrupts communication between these regions, weakening top-down executive control by the mPFC, while increasing reliance on habitual and subcortical processes that promote alcohol misuse (Abernathy et al., 2010; Nippert et al., 2024; Song et al., 2024). Consistent with this idea, manipulations of mPFC→NAcSh circuitry influence alcohol seeking, cue-induced reinstatement, and relapse-like behavior (Barker et al., 2015; Marchant et al., 2015; Moorman et al., 2015).

Despite compelling evidence implicating corticostriatal dysfunction in AUD, little is known regarding whether there are sex differences in the extent to which these projections guide drinking. Most mechanistic studies examining corticostriatal circuitry have been conducted in males animals (Beery & Zucker, 2011; Shansky & Woolley, 2016), despite growing evidence that males and females recruit partially distinct neural networks during alcohol dependence (Becker et al., 2012; Ardinger et al., 2024). Supporting this theory, we recently demonstrated that local field potential (LFP) activity recorded from the infralimbic mPFC (IL) and NAcSh accurately predict alcohol drinking and chronic intermittent alcohol vapor (CIA) exposure in males but not females, suggesting that corticostriatal communication may differentially contribute to alcohol-related behaviors across the sexes (Hewitt et al., 2025). However, because LFP recordings are inherently correlational, whether activity within IL-NAcSh pathway is functionally relevant in alcohol dependence, and whether this relevance is sex specific, remains unknown.

To address this gap, the present study utilized a projection-specific chemogenetic approach to manipulate IL→NAcSh neurons during acute withdrawal following CIA exposure. By selectively activating or inhibiting this pathway, we sought to determine whether NAcSh-projecting infralimbic mPFC (IL) neurons causally regulate alcohol self-administration and withdrawal-related anxiety-like behavior in males and females. The IL is particularly well positioned to regulate dependence-related phenotypes because it contributes to behavioral flexibility, extinction learning, inhibitory control, and the expression of goal-directed behaviors (Barker et al., 2015; Moorman et al., 2015; Peters et al., 2009). We therefore hypothesized that IL→NAcSh projections exert greater control over alcohol-related behaviors in males than in females. More specifically, we predicted chemogenetic manipulation of this pathway to produce weaker or no effect in females, consistent with evidence that this corticostriatal circuit is more relevant to male drinking behavior during alcohol withdrawal (Hewitt et al., 2025).

## METHODS

### Animals

Adult (approximately 70 days old) male and female Sprague-Dawley rats were used for these experiments. While Sprague-Dawley rats sometimes tend to drink less alcohol compared with other strains (Gauvin et al., 1993), Sprague-Dawleys are the most commonly used outbred strain in behavioral studies, and their high genetic diversity increases the translational relevance of this model (Gileta et al., 2022). Further, more recent data suggests that strain differences are not consistent across studies (Patwell et al., 2021; Sauton et al., 2021).

Rats were obtained from Inotiv (West Lafayette, IN, USA) and were housed in standard polycarbonate cages with *ad libitum* access to food and water. Rats were kept under a 12-hour reverse light-dark cycle (lights off at 08:00), allowing all experimental procedures to be conducted during the rats’ active phase. Additionally, experiments were performed at the same time every day to avoid potential circadian rhythm disruptions. Animal facilities were accredited by the American Association of Laboratory Animal Care (AALAC) and all procedures were approved by the Institutional Animal Care and Use Committee (IACUC) at Washington State University. Same-sex rats were housed 2-3 per cage prior to stereotaxic surgery, single-housed for 8 recovery days after surgery, and then reunited with their original cage mates. An overall experimental timeline can be found in **Figure 1**.

**Figure 1.**
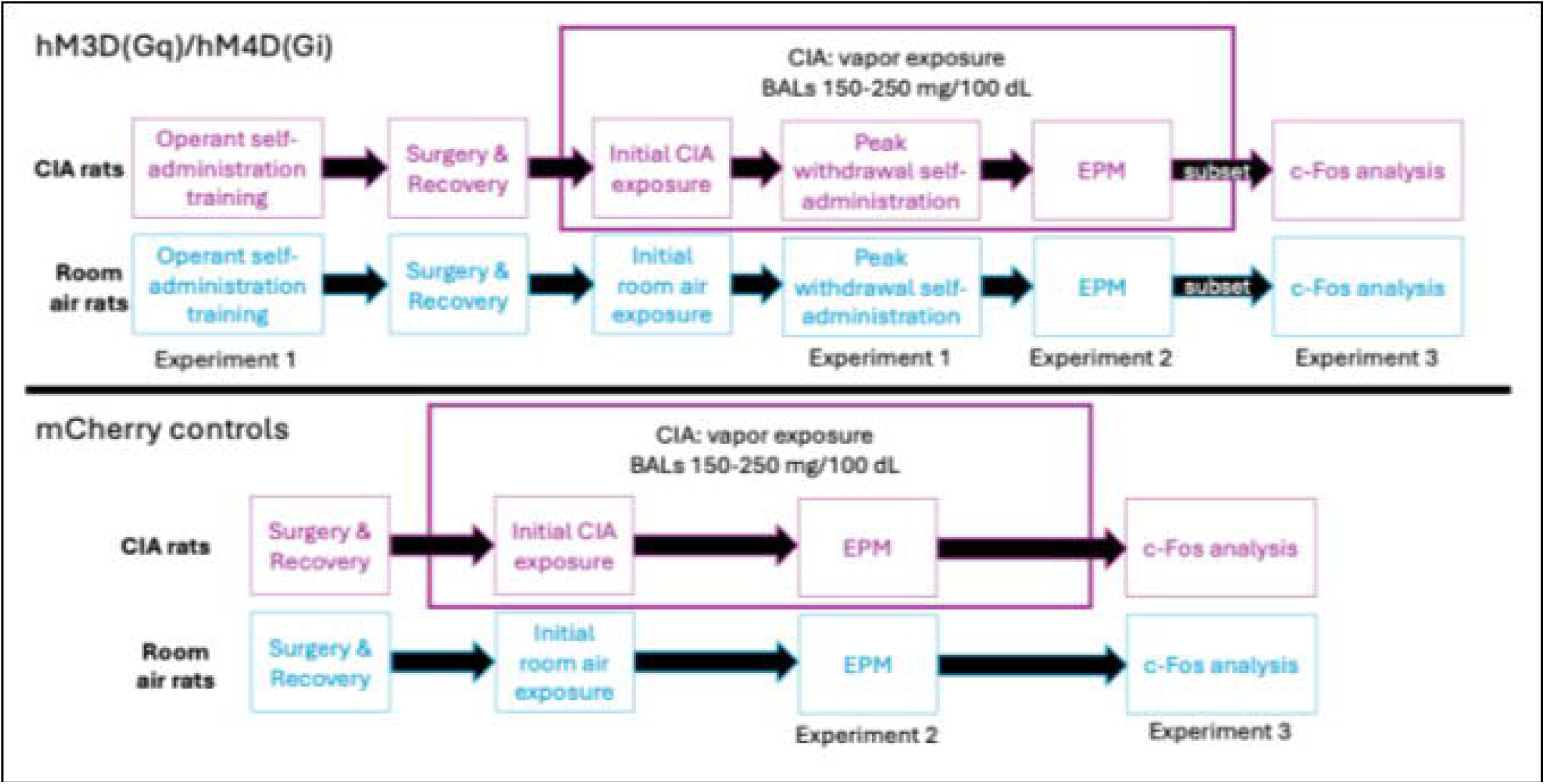
Experimental timeline.

### Manipulation of IL→NAcSh projections on alcohol self-administration during acute withdrawal

#### Alcohol self-administration training

Self-administration took place during the rats’ dark cycle in standard, sound attenuating operant boxes (10” L x 12.5” W x 8.5” H) programmed using the MED-Associated IV software. Boxes are equipped with two retractable levers on the same wall of the chamber with a center receptacle for liquid distribution. Presses to the active lever resulted in the delivery of 0.1mL of liquid. Responses on the inactive lever resulted in no reinforcer. A house light was illuminated for the duration that levers were available for each session (28.2 lux).

Rats were trained to self-administer 10% ethanol in water under fixed ratio-1 (FR-1) reinforcement schedule using a sucrose fade technique, as described in our previous work (Henricks et al., 2016; Hewitt et al., 2025; Nicholson et al., 2026). Rats were trained with the following sequence of solutions: 10% sucrose (4-5 sessions), 10% sucrose + 10% ethanol (5 sessions), 5% sucrose + 10% ethanol (5 sessions), 10% ethanol (15 sessions). Each session in the operant chambers lased 30 minutes and sessions were conducted 5 days per week, Monday – Friday. Based on our previous experience, training was considered complete after 15 sessions of 10% ethanol (Henricks et al., 2016; Hewitt et al., 2025; Nicholson et al., 2026). The average g/kg of alcohol consumed over the last four session was calculated for each animal as the pre-CIA drinking level.

#### Stereotaxic surgery

Rats then underwent stereotaxic surgery for DREADD injection. For all surgical procedures, anesthesia was induced by inhalation of 4-5% isoflurane (Med-Vet International, Mettawa, IL, USA) in an air-sealed chamber. Analgesia and antibiotics (meloxicam, 2 mg/kg, s.c.; enrofloxacin, 5 mg/kg, s.c., Med-Vet International, Mettawa, IL, USA) were administered after anesthesia induction and during the first three days of post-operative care. The surgical field was shaved and disinfected with ethanol and iodine antiseptic solution. OptixCare eye lubricant (Aventix Animal Health, Burlington, Ontario, CA) was applied to the eyes to prevent drying. All surgeries were performed under sterile conditions with the surgical plane determined by loss of pedal withdrawal and corneal reflexes. Once a deep level of anesthesia was obtained, rats were placed on a heating blanket and mounted into a stereotaxic frame. Rats received bilateral 0.3 µL injections of either a cre-dependent excitatory (pAAV-hSyn-DIO-hM3D(Gq)-mCherry (AAV5); Addgene, catalog #44361-AAV5, titer 6 x 10^12^ vg/ml), inhibitory (pAAV-hSyn-DIO-hM4D(Gi)-mCherry (AAV5); Addgene catalog #44362-AAV5, titer 6 x 10^12^ vg/ml), or mCherry control virus (pAAV-hSyn-DIO-mCherry (AAV5); Addgene catalog #50459-AAV5, titer 6 x 10^12^ vg/ml) into the IL mPFC (from bregma: DV -4.5mm, AP +3.0mm, ML +/-0.7mm). In the same surgery, rats were also bilaterally infused with 0.5 µL of a retrograde viral vector containing Cre recombinase (pENN.AAV.hSyn.HI.eGFP-Cre.WPRE.SV40 (AAV retrograde); Addgene catalog #105540-AAVrg, titer 7 x 10^12^ vg/ml) into the NAcSh (from bregma: DV –7.0 mm, AP +1.2 mm, ML +/-1.0 mm). All injections were infused at a rate of 150 µL/minute using a 10 µL Gastight Neuros Syringe (Hamilton Company catalog # 65460-06, Reno, NV, USA) and microinjection syringe pump (World Precision Instruments catalog # UMP3T-1, Sarasota, FL, USA). The needle was kept in place for an additional ten minutes to allow for diffusion before the syringe was slowly removed. Three-to-five silk sutures were used to close the wound. This intersectional chemogenetic approach enables selective manipulation of IL neurons that project to the NAcSh. A schematic of the chemogenetic approach is in **Figure 2**. Histological verification of viral injection sites in the IL and NAcSh is shown in **Figure 3**.

**Figure 2.**
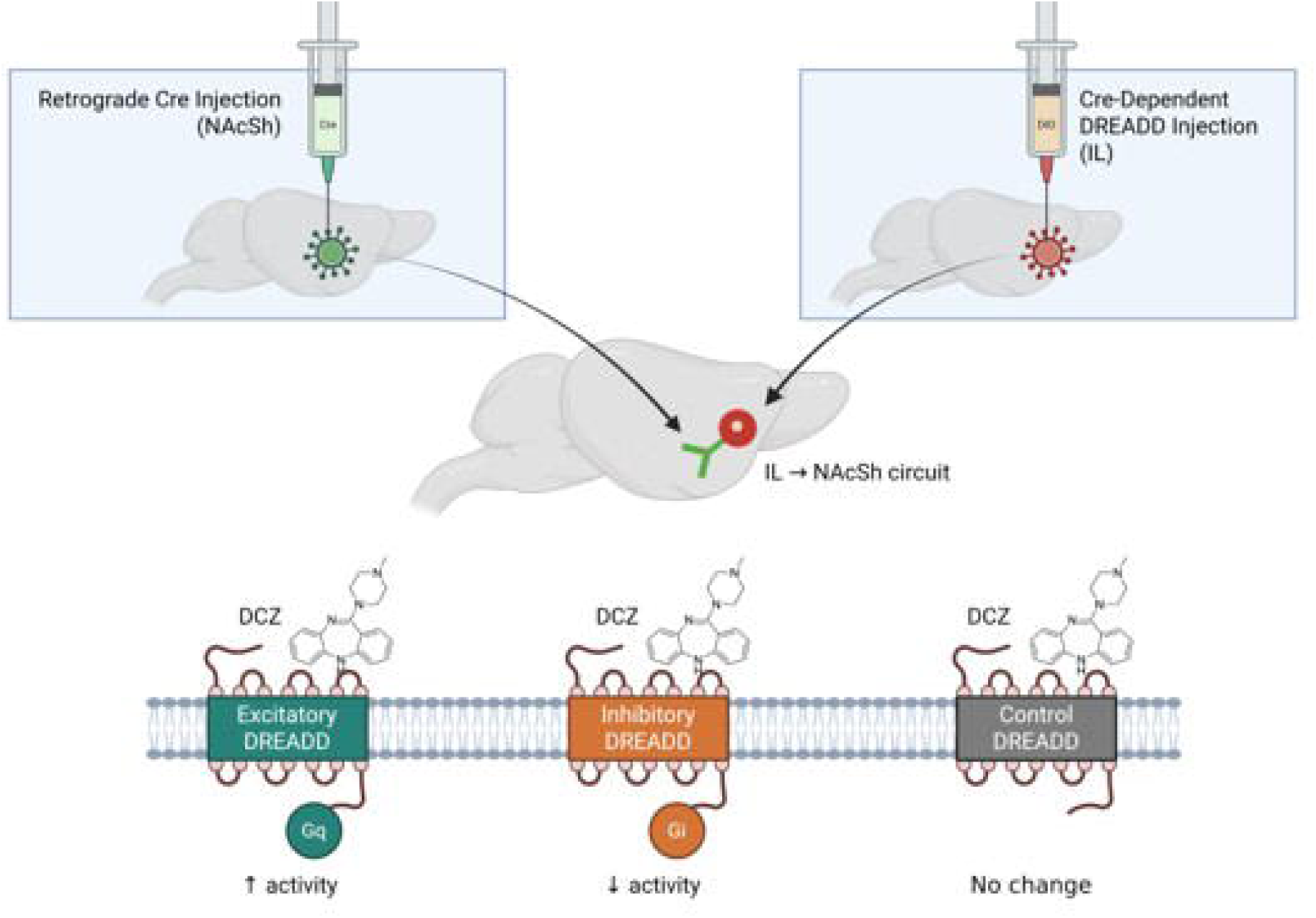
Conceptual schematic of DREADD viral strategy. Projection-specific DREADD strategy targeting IL projections to the NAcSh. A retrograde viral vector expressing Cre-recombinase was injected into the NAcSh, where it was taken up by axon terminals and transported back to the cell bodies of neurons projecting to the NAcSh, resulting in GFP expression (green). A Cre-dependent excitatory (hM3D(Gq), inhibitory (hM4D(Gi), or inactive control DREADD virus was injected into the IL, producing mCherry fluorescence (red). Consequently, only IL neurons projecting to the NAcSh co-expressed Cre and the DREADD receptor. Systemic administration of DCZ selectively activated or inhibited the DREADD receptors in these projection-defined neurons, enabling top-down manipulation of the ILà NAcSh pathway.

**Figure 3.**
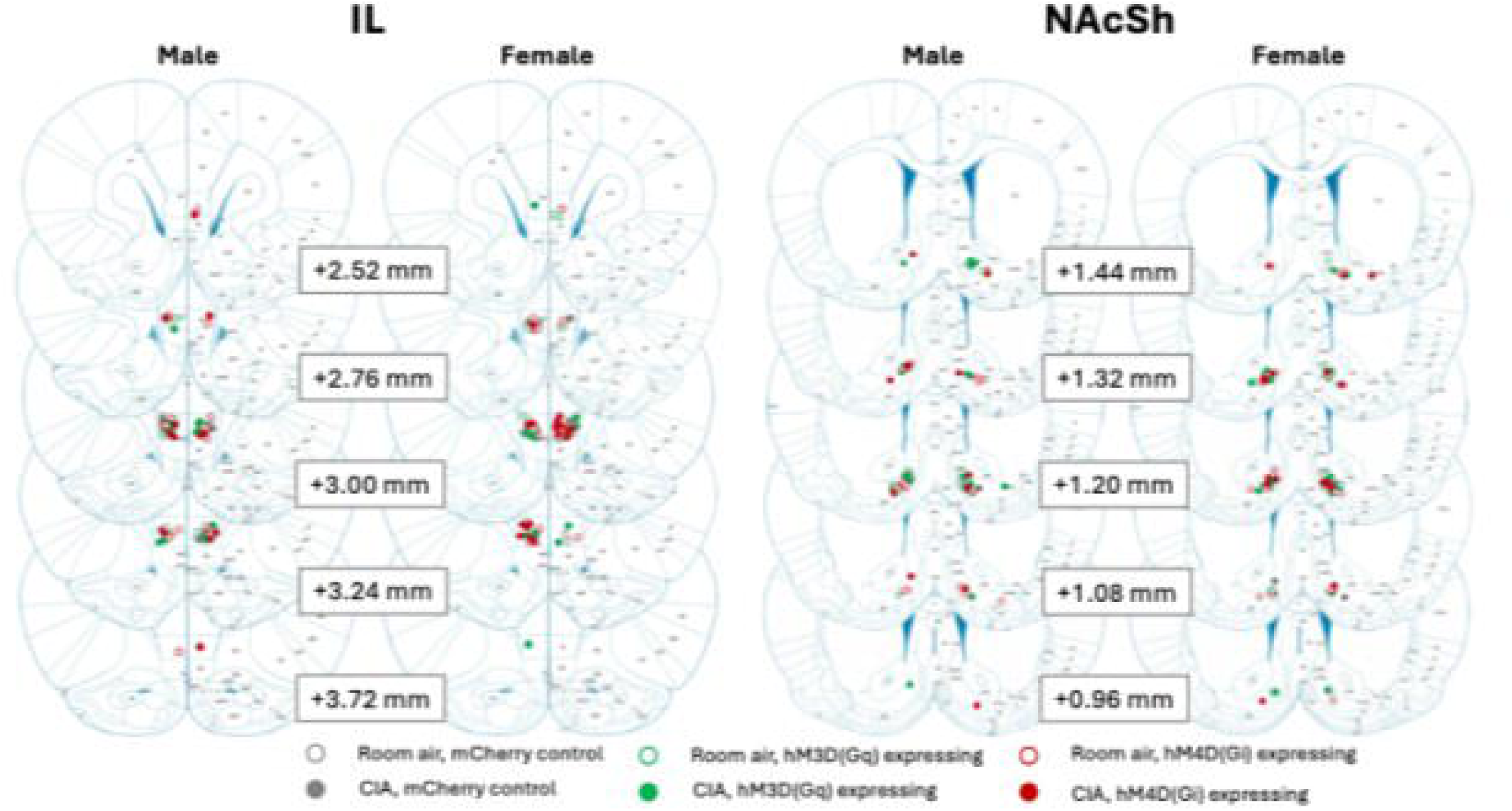
Histological verification of IL and NAcSh injection sites. IL slices are on the left and NAcSh is shown on the right which are further broken down with males on the left and females on the right. Closed circles indicate chronic intermittent alcohol (CIA) rats, while open circles indicate control rats. Green circles indicate hM3D(Gq)-expressing rats, red circles indicate hM4D(Gi)-expressing rats, and gray circles indicate rats who received the mCherry control DREADD. Histology was based on Paxinos & Watson: The rat brain in stereotaxic coordinates, 5^th^ edition.

After completion of the surgery, rats received 1.0 mL of 0.9% sterile saline s.c. to prevent dehydration and were placed in a clean cage on a heating pad for initial recovery. Rats received post-operative care for the following three days and then continued to be monitored daily for an additional five days to assess coat condition, body weight, general responsiveness/activity, and wound healing for any signs of infection. Once deemed fully healed and sutures were removed, rats were reunited with their original cage mates and allowed to continue to the next phase of the experiment.

#### Chronic intermittent alcohol exposure

After recovery, rats were either exposed to CIA vapor (*n* = 22 males, 24 females) or normal room air (control; *n* = 22 males, 22 females) for 6 weeks. Our custom-build, four-chamber apparatus (La Jolla Alcohol Research Inc., La Jolla, CA, USA) allows for blood alcohol concentrations (BACs) to be titrated by the experimenter by adjusting the rate of 95% ethanol that is vaporized and integrated into the air flow supplying the sealed Plexiglas boxes (25.5” L x 21” W x 13” H) in which the rats’ home cages were housed. CIA rats were exposed to vaporized ethanol for 14 hours per day from approximately 18:00-08:00; for 10 hours a day, they were just exposed to room air that was filtered into the chamber. Exposing rats to alcohol vapor predominately during their light phase allowed for testing acute withdrawal behavior during their active (dark) phase.

Rats in the CIA condition remained in the vapor chambers for the entirety of the experiment. Approximately 0.1 mL of blood was collected from the tail vein a minimum of two times per week and on testing days, right as the vapor turned off to ensure BACs remained at an appropriate level (150-250mg%), as described previously (Williams et al., 2012; Henricks et al., 2016; Hewitt et al., 2025). Bloods were centrifuged at 10,000 rpm at 4 °C to isolate plasma. Plasma samples (5μL) were then tested using an Analox AM 1 or Analox GL 5 (Analox Instruments Ltd., Lunenburg, MA, USA) for quantification of BACs.

#### Alcohol self-administration during acute withdrawal

After four weeks of CIA vapor or room air exposure, rats resumed alcohol self-administration for eight baseline sessions while their assigned exposure continued. In CIA-exposed rats, sessions took place during acute withdrawal, approximately 7–8 h after vapor turned off. After baseline testing, rats completed the vehicle session followed by the DCZ session, with one intervening day. By the DCZ session, rats had undergone approximately six weeks of CIA vapor or room air exposure. CIA exposed rats had to meet the criteria of escalating their self-administration rates to 1.5 times their pre-CIA levels in order to be included in the study.

#### Chemogenetic manipulation

After the eight baseline self-administration sessions, rats were pretreated with 0.1 mg/kg of the DREADD agonist deschloroclozapine (DCZ) or vehicle (DMSO into sterile saline) 30 minutes before the beginning of the session. DCZ (HelloBio, catalog # HB9126) was dissolved in DMSO at a concentration of 25 mg/ml. This was further diluted into 0.9% sterile saline for a concentration of 0.1 mg/ml. In order to allow for a within-subjects design for the self-administration experiments, all rats received vehicle pretreatment followed by DCZ pretreatment, with one day between these sessions in order to prevent a carry-over effect of DREADD activation.

### Manipulation of IL→NAcSh projections on anxiety-like behavior in acute withdrawal

#### Elevated plus maze

After self-administration testing was complete, rats underwent elevated plus maze (EPM) testing during acute withdrawal. Because the EPM does not allow for repeated testing in a short time-frame due to habituation (Schrader et al., 2018), we used a between-subject design for this experiment. An independent cohort of rats expressing mCherry control vectors was included in EPM testing, but were not trained to self-administer alcohol. However, the mCherry control cohort was also exposed to 6 weeks of CIA or room air exposure in order to be time-matched to the active DREADD cohorts. All rats received a 30-minute pretreatment of 0.1 mg/kg DCZ prior to testing.

Rats were placed in the center of the maze and allowed to freely explore the apparatus in dim lighting for 5 minutes (Open arm = 45.6 lux; Closed arm = 2.0 lux; EPM center = 20.3 lux). Sessions were recorded and the time spent in closed and open arms (seconds), as well as distance traveled (cm), was analyzed using EthoVision XT (Noldus, Leesberg, VA, USA). A subset of videos was manually scored by blinded reviewers to verify the accuracy of the automated tracking software.

### Manipulation of IL→NAcSh projections on IL c-Fos activation

#### Tissue collection

Within a week of cessation of behavioral testing, a portion of rats received a 120-minute pretreatment of DCZ before an overdose with isoflurane and were immediately perfused with ice-cold 1x phosphate buffered saline (1x PBS), followed by 4% paraformaldehyde dissolved in 1x PBS (PFA). This pretreatment time allowed us to capture DCZ-induced changes in c-Fos expression at what would be the approximate beginning of the self-administration sessions. Brains were post-fixed overnight in PFA at 4 °C and then placed into 30% sucrose until they sank. They were then rapidly frozen in powdered dry ice and stored at −80 °C. Brains were sectioned coronally at the IL injection site (40 μm thickness) at -20 °C using a cryostat (Leica Biosystems, Buffalo Grove, IL, USA). Sectioned tissue was stored at 4°C in 0.1% sodium azide dissolved in 1x PBS.

#### Immunohistochemistry

Tissue sections were first washed three times in 1x PBS + 0.1% Triton-X, then washed in blocking buffer (1x PBX + 0.1% Triton-X + 3% Normal Donkey Serum) for 1-2 hours, and then incubated overnight at room temperature in blocking buffer and primary antibodies (rabbit anti-c-Fos, 1:1000 dilution, Synaptic Systems, catalog # 226 008; chicken anti-GFP, 1:1000 dilution, Abcam, catalog # ab13970; Guinea pig anti-mCherry, 1:10,000 dilution, Synaptic Systems, catalog # 390-004). Following primary incubation, tissue was washed 3 times in 1X PBS and 0.1% Triton-X for 10 minutes each and then incubated for 2-4 hours in blocking buffer and secondary antibodies from Jackson ImmunoResearch Laboratories (Alexa Fluor® 647 AffiniPure® Donkey Anti-Rabbit IgG (H+L), catalog # 711-605-152; Alexa Fluor® 488 AffiniPure® Donkey Anti-Chicken IgY (IgG) (H+L), catalog # 703-545-155; Cy™3 AffiniPure® Donkey Anti-Guinea Pig IgG (H+L), catalog # 706-165-148) all diluted 1:500). Tissue was then washed three times in 1x PBS before a five-minute incubation in DAPI (Invitrogen, catalog # P36931) diluted 1:1000 in 1x PBS. Tissue sections were then washed another three times in 1x PBS and immediately mounted on standard, positively charged microscope slides (Globe Scientific) and cover-slipped using ProLong Gold anti-fade mounting media (Thermo Fisher, catalog # P10144).

#### Quantification of c-Fos-positive cells

The brain region of interest (IL) was identified using DAPI staining and the Paxinos & Watson rat brain atlas. Mounted tissue was imaged using an epifluorescence microscope (Leica CTR 6500, Wetzler, Germany). The 405, 488, 561, and 640 nm lasers were used to obtain images of DAPI, GFP, mCherry, and c-Fos respectively. Image acquisition settings (e.g., laser power settings, gain, zoom) were held constant across all images to enable proper comparison of c-Fos induction slices expressing or lacking DREADD expression. Two images were collected from the IL for each brain. Exported image files were analyzed in ImageJ to quantify the number of c-Fos+, GFP+, and mCherry+ colocalization. Cell counts were performed independently by two individuals blind to experimental condition. Counts between the two raters were averaged unless they differed by more than a standard deviation of ±2, in which case a third blinded reviewer re-examined the image and determined the final count. c-Fos immunoreactivity was quantified as the percentage of DAPI-positive nuclei that were c-Fos positive within the IL region of interest. Representative c-Fos images from the IL are shown in **Figure 4**.

**Figure 4.**
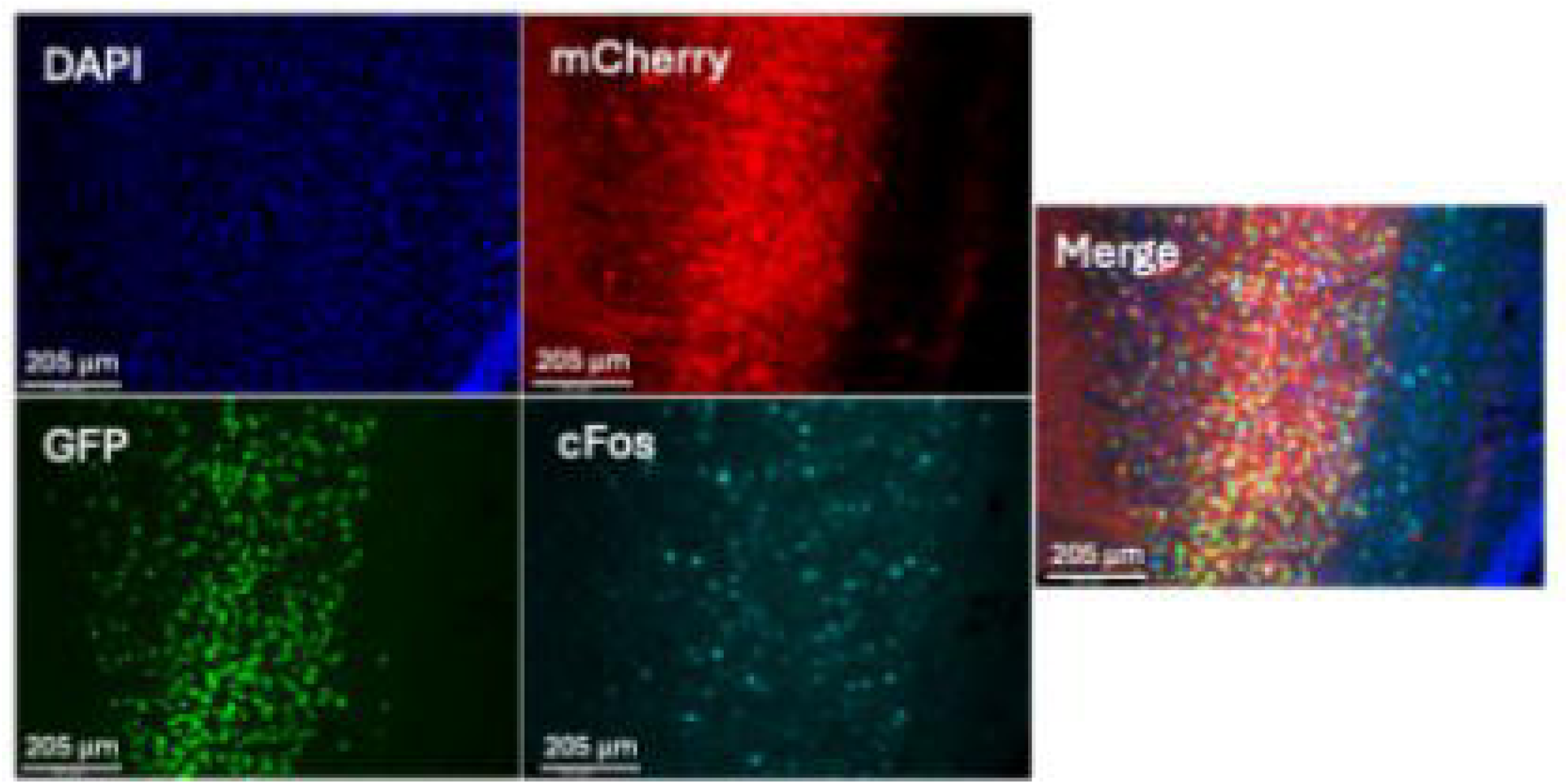
Representative images of the IL showing immunohistochemical fluorescence for DAPI (blue), mCherry (DREADD-red), GFP (retrograde cre recombinase - green), c-Fos activation (cyan), and a merged image at 10x magnification.

### Statistical analyses

Five rats that had misplaced, or not visible (due to microinjector malfunction) stereotaxic injections were excluded from all data analysis along with three CIA exposed rats who did not meet the escalation criteria (escalation of 1.5x baseline self-administration average). Male and female data were analyzed separately. To determine how alcohol self-administration was impacted by CIA and DREADD manipulation, mixed factorial ANOVAs were conducted with CIA exposure (room air control, CIA) and DREADD type (hM3D(Gq), hM4D(Gi)) as between-subjects factors. The average of the last four training sessions were quantified as the “Pre-CIA” baseline, and the average of the last four sessions during CIA-induced withdrawal, prior to vehicle pretreatment, were averaged as the “CIA baseline.” Thus, the within-subjects factor was session type (pre-CIA, CIA baseline, vehicle pretreatment, and DCZ pretreatment). G/kg consumed and number of inactive lever presses served as dependent variables.

To determine how anxiety-like behavior in the EPM and c-Fos immunoreactivity were impacted by CIA and DREADD manipulation, 2x3 ANOVAs were used with CIA exposure (room air, CIA) and DREADD type (mCherry control, hM3D(Gq), hM4D(Gi)) as the between-subjects factors. For the EPM, time (s) spent in the open and closed arms, and locomotor activity (distance traveled; cm) were the dependent variables. The number of c-Fos+, mCherry and GFP expressing cells was quantified, and were analyzed as the percentage of c-Fos+ cells relative the total number of cells in the region.

## RESULTS

### Manipulation of IL→NAcSh projections on alcohol self-administration during acute withdrawal

#### G/kg of alcohol consumed

For males, a mixed factorial ANOVA revealed a significant main effect of session [*F*(3,87) = 22.80, *p* < .001, = .44], CIA [*F*(1,29) = 21.03, *p* < .001, = .42], a session*CIA interaction [*F*(3,87) = 9.26, *p* < .001, = .24], and a session*CIA*DREADD interaction [*F*(3,87) = 3.72, *p* = .01, = .11] on g/kg of alcohol consumed (n = 7-10/group; **Figure 5 A & B**). There was no main effect of DREADD type [*F*(1,29) = .12, *p* = .73, = .004] or any other significant interactions (*p*s > .05). For the session*CIA*DREADD interaction, post-hoc t-tests indicated that control males expressing hM3D(Gq) consumed more alcohol (g/kg) after DCZ than at CIA Baseline or after vehicle (*ps* < .05). In CIA males, alcohol consumed in CIA Baseline, Vehicle and DCZ sessions were all significantly higher than Pre-CIA levels regardless of DREADD type (*ps* < .05). Importantly, hM4D(Gi) manipulation by DCZ selectively increased g/kg of alcohol consumed compared to vehicle treatment (*p* < .001).

**Figure 5.**
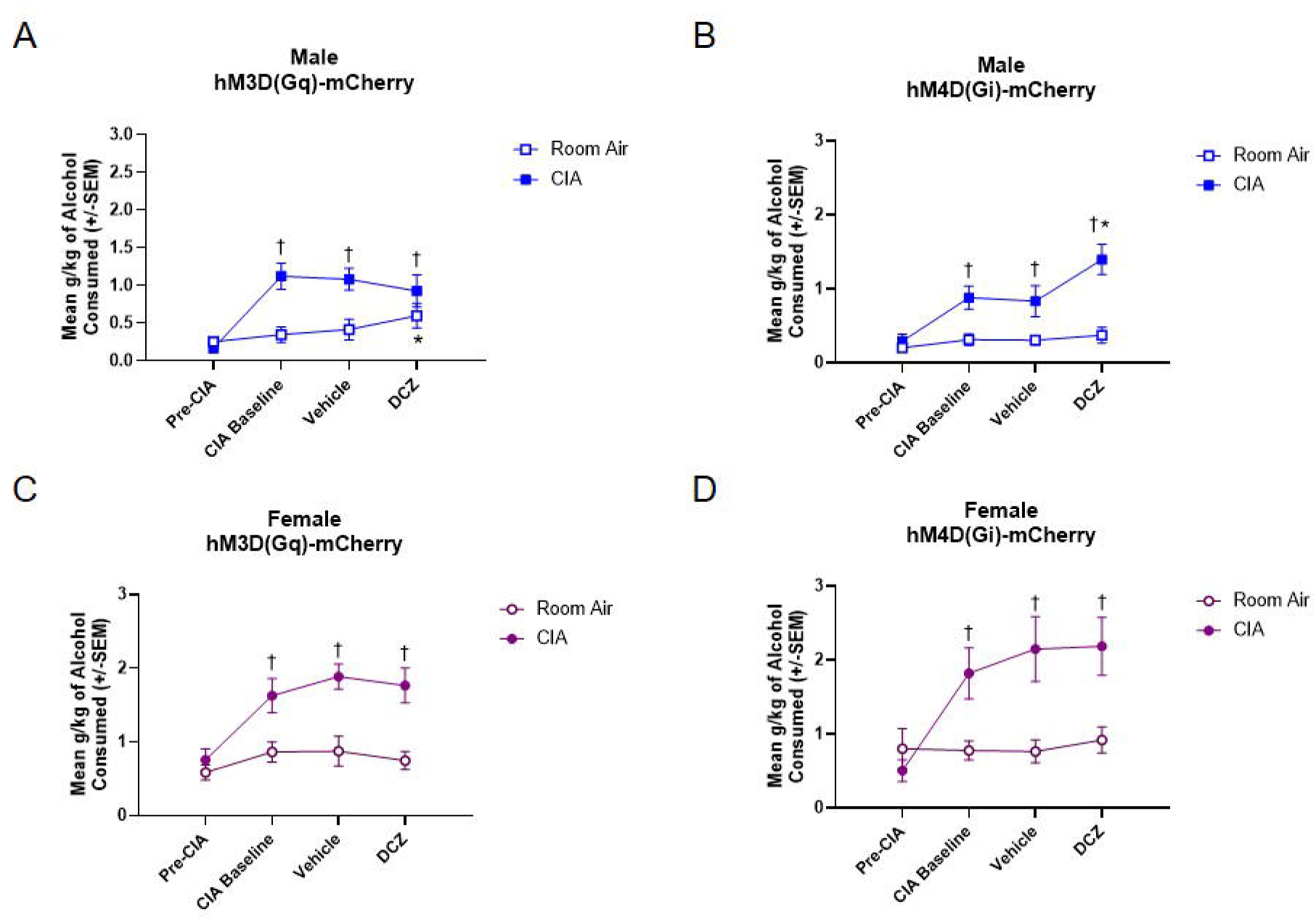
Alcohol self-administration in male and female rats following chronic intermittent alcohol exposure. Alcohol intake (g/kg of body weight) during 30-min self-administration sessions is shown for male (A, B) and female (C, D) rats exposed to CIA or room air (n = 7-10/group/sex). Rats expressed hM3D(Gq) (A, C) or hM4D(Gi) (B, D). Intake was measured before the 6-week exposure period (Pre-CIA), and during acute withdrawal after the exposure period (CIA Baseline), and in sessions preceded by a 30-min pretreatment with vehicle or DCZ. ^†^In CIA-exposed males and females, intake was higher at CIA Baseline, Vehicle, and DCZ than at Pre-CIA, across DREADD types (*ps* < .05). *In room air control males expressing hM3D(Gq) (A), intake was higher after DCZ than at CIA Baseline and after vehicle (*ps* < .05). *In CIA-exposed males expressing hM4D(Gi) (B), intake was higher after DCZ than after vehicle (*p* < .001).

In females, a mixed factorial ANOVA revealed a significant main effect of session [*F*(3,90) = 17.46, *p* < .001, = .37], CIA [*F*(1,30) = 16.40, *p* < .001, = .35], and a session*CIA interaction [*F*(3,90) = 11.69, *p* < .001, = .28] on g/kg of alcohol consumed (n = 7-9/group; **Figure 5 C & D**). There was no main effect of DREADD type [*F*(1,30) = .28, *p* = .60, = .01] or any other significant interactions (*p*s > .05). In CIA females, alcohol consumed in CIA Baseline, Vehicle and DCZ sessions were all significantly higher than Pre-CIA levels regardless of DREADD type (*ps* < .05)

#### Inactive lever presses

For males, a mixed factorial ANOVA revealed a significant main effect of session [*F*(3,87) = 3.10, *p* = .03, = .10] on inactive lever presses (**Figure 6 A & B**). There was no main effect of CIA [*F*(1,29) = 2.86, *p* = .10, = .10] or DREADD type [*F*(1,29) = 1.00, *p* = .33, = .03], nor any other significant interactions (*ps* > .05).

**Figure 6.**
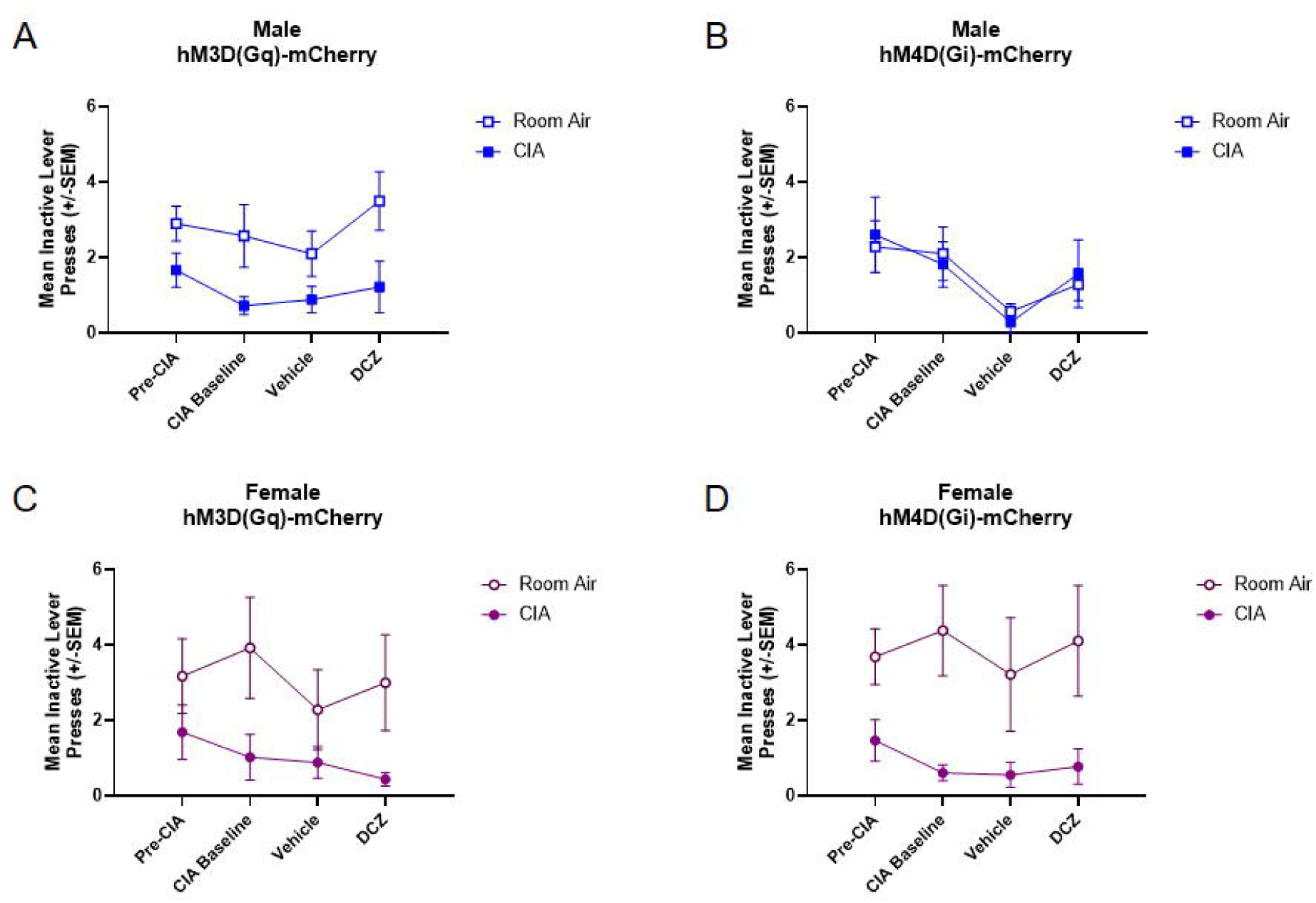
Inactive lever presses during alcohol self-administration. Inactive lever presses during 30-min self-administration sessions are shown for male (A, B) and female (C, D) rats exposed to CIA or room air (n = 7-10/group/sex). Rats expressed hM3D(Gq) (A, C) or hM4D(Gi) (B, D). Lever presses were measured before the 6-week exposure period (Pre-CIA), and during acute withdrawal after the exposure period (CIA Baseline), and in sessions preceded by a 30-min pretreatment with vehicle or DCZ. Mixed factorial ANOVAs revealed a main effect of session in males (*p* = .03), and a main effect of CIA exposure in females (*p* < .001).

In females, a mixed factorial ANOVA revealed a significant main effect of CIA [*F*(1,30) = 15.82, *p* < .001, = .35] on inactive lever presses (**Figure 6 C & D**). There was no main effect of session [*F*(3,90) = 1.08, *p* = .36, = .04] or DREADD type [*F*(1,30) = .24, *p* = .63, = .01], nor any other significant interactions (*ps* > .05).

### Manipulation of IL→NAcSh projections on anxiety-like behavior in acute withdrawal

#### Time spent in open and closed arms

For males, a 2x3 ANOVA revealed a significant main effect of DREADD type [*F*(2, 38) = 6.52, *p* = .004, = .26] on time spent in the open arm of the EPM (n = 5-10/group; **Figure 7A**). There was no main effect of CIA [*F*(1, 38) = 0.25, *p* = .62, = .01] or a significant interaction (*p* > .05). There was also a significant main effect of DREADD type [*F*(2, 38) = 4.82, *p* = .01, = .20] for time spent in the closed arm of the EPM, but no effect of CIA [*F*(1, 38) = 0.29, *p* = .59, = .008] or a significant interaction (*p* > .05; **Figure 7B**). Further interrogation of the DREADD effects showed that hM3D(Gq) manipulation by DCZ reduced open arm time and increased closed arm time in the EPM relative to mCherry controls (*ps* < .05).

**Figure 7.**
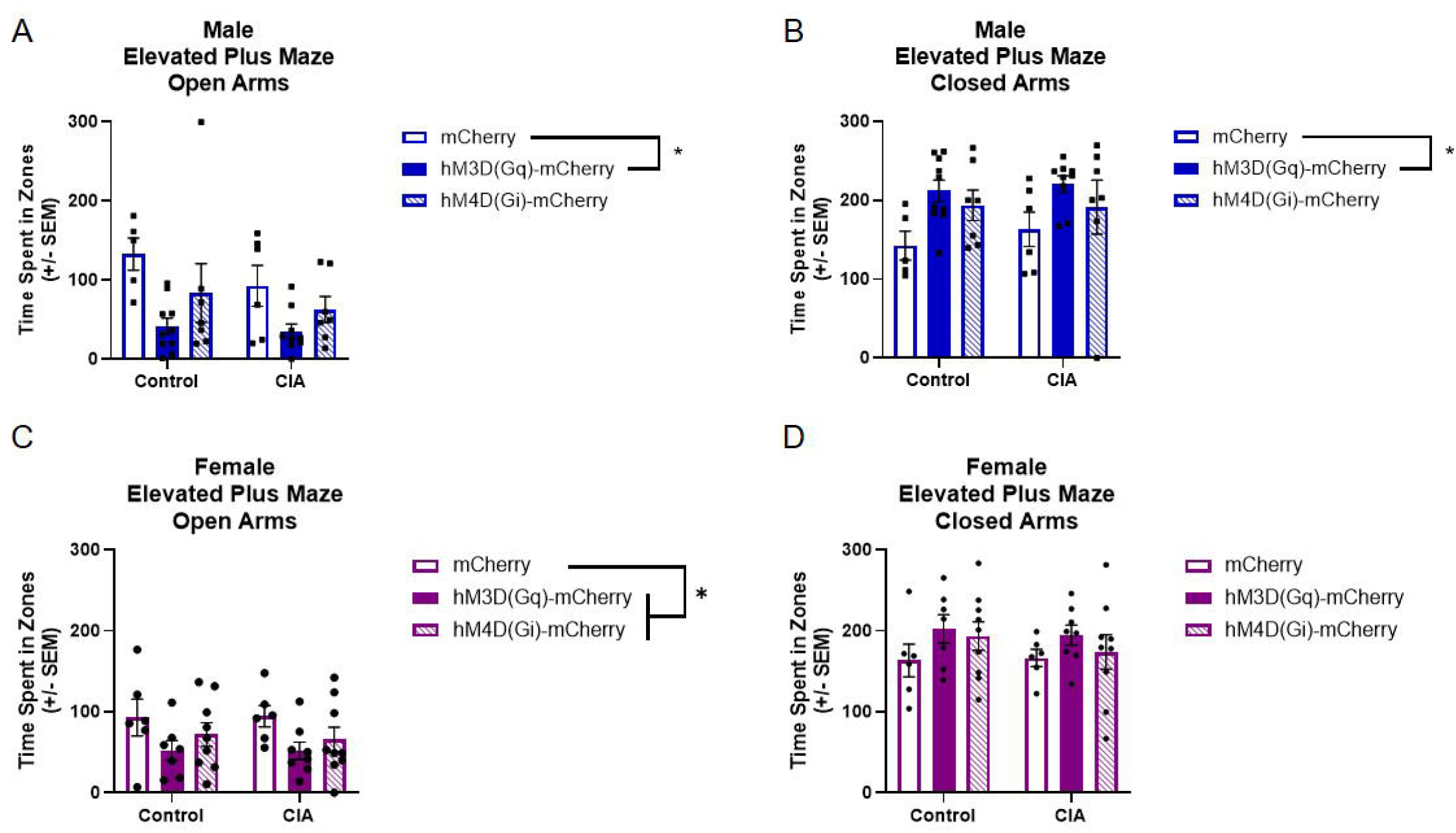
Time spent in the open and closed arms of the elevated plus maze following DCZ administration. Open-arm time is shown for males (A) and females (C), and closed-arm time is shown for males (B) and females (D). Rats were exposed to room air or CIA and expressed mCherry, hM3D(Gq), or hM4D(Gi) (*n* = 5–10 per group for males and 6–9 per group for females). *Males expressing hM3D(Gq) spent less time in the open arms (A) and more time in the closed arms (B) than mCherry controls (*ps* < .05). *Females expressing hM3D(Gq) or hM4D(Gi) spent less time in the open arms (C) than mCherry controls (*ps* < .01).

In females, a 2x3 ANOVA revealed a significant main effect of DREADD type [*F*(2, 29) = 4.22, *p* = .03, = .23] on time spent in the open arm of the EPM (n = 6-9/group; **Figure 7C**). There were no main effect of CIA [*F*(1, 29) = .21, *p* = .65, = .01] or a significant interaction (*p* > .05). Further interrogation of the DREADD effect showed that both hM3D(Gq) and hM4D(Gi) manipulation by DCZ reduced open arm time in the EPM relative to mCherry controls (*ps* < .01). There were no significant main effects of DREADD type [*F*(2, 29) = 1.77, *p* = .19, = .11] or CIA [*F*(1, 29) = .05, *p* = .82, = .002] for time spent in the closed arm of the EPM, or a significant interaction (*p* > .05; **Figure 7D**).

#### Distance traveled

For males, a 2x3 ANOVA revealed a significant main effect of DREADD type [*F*(2, 38) =12.83, *p* < .001, = .40] and a CIA*DREADD interaction [*F*(2, 38) = 3.84, *p* = .03, = .17] on distance traveled (cm) in the EPM (n = 5-10/group; **Figure 8A**). There was no main effect of CIA [*F*(1, 38) = 2.48, *p* = .12, = .06]. Post-hoc t-tests suggest that both hM3D(Gq) and hM4D(Gi) manipulation by DCZ reduced distance traveled in the CIA males compared to the mCherry controls (*ps* < .05).

**Figure 8.**
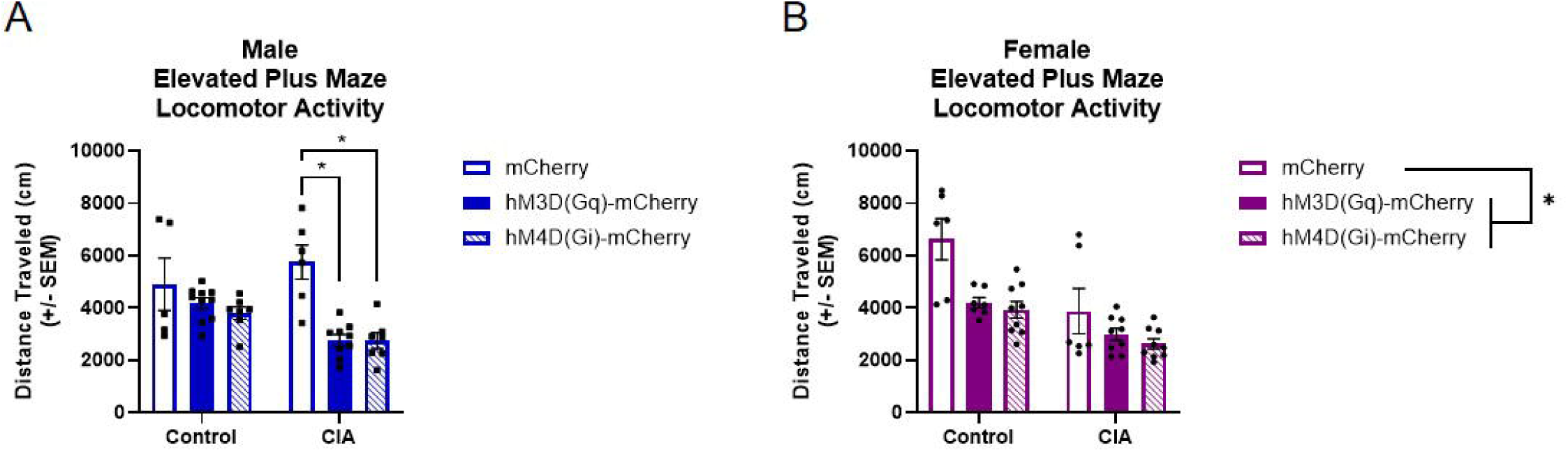
Distance traveled in the elevated plus maze following DCZ administration. Distance traveled (cm) is shown for male (A) and female (B) rats exposed to room air or CIA and expressing mCherry, hM3D(Gq), or hM4D(Gi) (*n* = 5–10 per group for males and 6–9 per group for females). *In males (A), CIA-exposed rats expressing hM3D(Gq) or hM4D(Gi) traveled less than CIA-exposed mCherry controls (*ps* < .05). *In females (B), rats expressing hM3D(Gq) or hM4D(Gi) traveled less than mCherry controls, across exposure conditions (*ps* < .05). In general, CIA-exposed females traveled less than room-air controls (main effect of CIA, *p* < .001).

In females, a 2x3 ANOVA revealed a significant main effect of DREADD type [*F*(2, 29) = 6.07, *p* = .006, = .30] and CIA [*F*(1, 29) =15.41, *p* < .001, = .35], but no interaction (*p* > .05; n = 6-9/group; **Figure 8B**). Further interrogation of the DREADD effects suggest that both hM3D(Gq) and hM4D(Gi) manipulation by DCZ reduced distance traveled in the EPM relative to mCherry controls (*ps* < .05).

### Manipulation of IL→NAcSh projections on IL c-Fos expression

For males, a 2x3 ANOVA revealed a significant main effect of CIA [*F*(1, 29) = 9.73, *p* = .004, = .25], DREADD type [*F*(2, 29) = 200.16, *p* < .001, = .93] and a CIA*DREADD interaction [*F*(2, 29) = 4.28, *p* = .02, = .23] on the percentage of c-Fos+ cells in the IL (n = 5-7/group; **Figure 9A**). Post-hoc t-tests showed that, within each exposure condition, the percentage of c-Fos+ cells in the IL was highest in hM3D(Gq) rats, intermediate in mCherry controls, and lowest in hM4D(Gi) rats (*ps* < .05).

**Figure 9.**
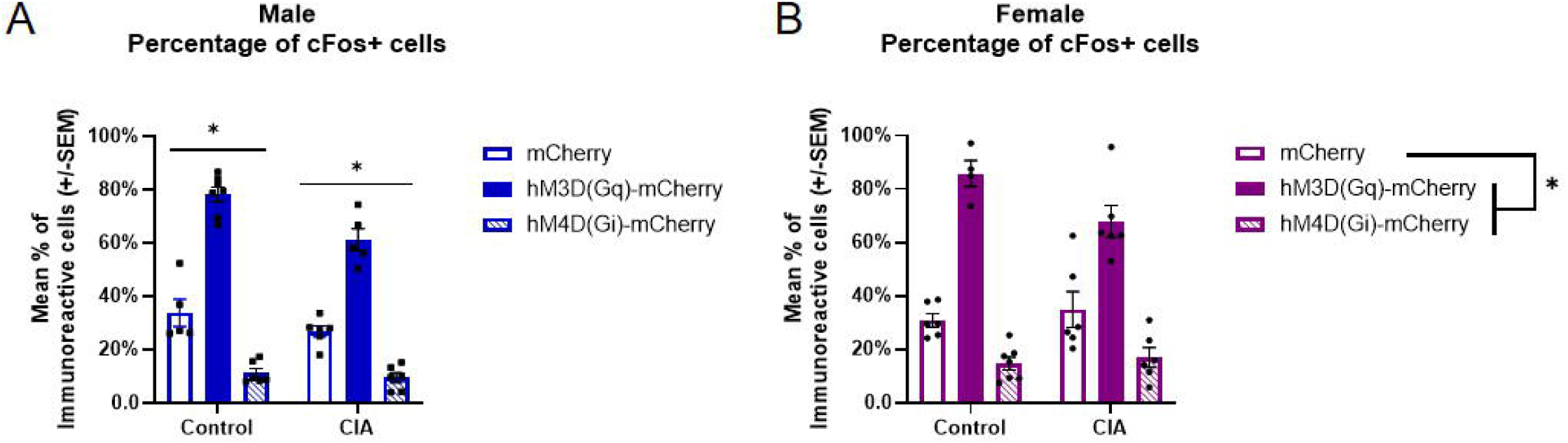
c-Fos immunoreactivity in the infralimbic cortex following DCZ administration. The percentage of c-Fos+ cells in the IL is shown for male (A) and female (B) rats exposed to room air or CIA and expressing mCherry, hM3D(Gq), or hM4D(Gi) (*n* = 5–7 per group for males and 4–7 per group for females). *In males (A), the percentage of c-Fos+ cells was highest in hM3D(Gq) rats, intermediate in mCherry controls, and lowest in hM4D(Gi) rats within each exposure condition (*ps* < .05). *In females (B), the percentage of c-Fos+ cells was higher in hM3D(Gq) rats and lower in hM4D(Gi) rats relative to mCherry controls, across exposure conditions (*ps* < .05).

In females, a 2x3 ANOVA revealed a significant main effect of DREADD type [*F*(2, 29) = 86.43, *p* < .001, = .86] on the percentage of c-Fos+ cells in the IL (n = 4-7/group; **Figure 9B**). There was no effect of CIA [*F*(1, 29) = 2.97, *p* = .10, = .09] or a significant interaction (*p* > .05). Further interrogation of the DREADD effects suggest that hM3D(Gq) manipulation by DCZ increased c-Fos immunoreactivity, while hM4D(Gi) manipulation by DCZ reduced c-Fos immunoreactivity relative to mCherry controls, regardless of CIA exposure (*ps* < .05).

## DISCUSSION

The present study investigated whether NAcSh-projecting IL neurons control alcohol self-administration and anxiety-like behavior in alcohol dependent male and female rats. Chemogenetic inhibition of the IL→NAcSh pathway selectively increases alcohol self-administration in CIA-exposed males, supporting the hypothesis that this corticostriatal projection helps constrain alcohol intake during acute withdrawal in male rats. In room air control males, activation increased intake, suggesting that the behavioral consequences of manipulating this circuit depend on alcohol exposure history. In contrast, alcohol self-administration in females remained unaffected by IL→NAcSh manipulation, despite escalation after CIA exposure and changes in IL c-Fos immunoreactivity in the expected directions in both sexes. The effects on EPM behavior diverged from those on drinking. For example, both manipulations reduced open-arm time in females without detectably changing alcohol intake. This divergence suggests that the influence of IL→NAcSh neurons on anxiety-related behavior and alcohol self-administration may be partly dissociable, raising the possibility that these neurons constrain withdrawal-associated drinking through mechanisms beyond their influence on negative affect. Together, these findings extend our previous correlational LFP work by demonstrating a causal role for IL→NAcSh circuitry in alcohol dependence in males, and support the broader concept that males and females might achieve similar alcohol-related behaviors through different circuit strategies.

While chemogenetic inhibition of the NAcSh-projecting IL neurons selectively increased self-administration in alcohol-dependent males, excitation not only failed to reduce drinking, but slightly enhanced drinking in control males. These findings indicate that endogenous activity within the IL→NAcSh pathway is necessary for constraining excessive alcohol consumption in dependent males, but is insufficient on its own to reverse dependence-like drinking behavior. Thus, rather than functioning as an independent regulator of alcohol intake, this corticostriatal projection appears to represent one component of a broader neural network that becomes dysregulated following alcohol exposure. This interpretation is consistent with models of addiction proposing that dependence emerges through widespread neuroadaptations across corticostriatal, limbic, and stress-related circuitry rather than dysfunction within a single pathway (Koob, 2014; Koob & Volkow, 2016).

Although excitation of the IL→NAcSh projection did not reduce alcohol self-administration in this study in either sex, this finding should not be interpreted as evidence that increased activity within this pathway is behaviorally irrelevant during alcohol dependence. Rather, contemporary evidence suggests that IL→NAcSh signaling primarily contributes to behavioral inhibition and flexible updating of learned behaviors rather than the regulation of ongoing reward consumption. Consistent with this interpretation, chemogenetic activation of ventromedial PFC projections to the NAcSh suppresses cue-induced reinstatement of cocaine seeking after extinction, without affecting cocaine self-administration in animals who had not undergone extinction training (Augur et al., 2016). Because rats in the present study were actively self-administering alcohol during stable reinforcement rather than undergoing extinction or reinstatement, chemogenetic excitation may have occurred outside of the behavioral context in which this pathway normally exerts the greatest influence. Future studies should determine whether IL→NAcSh activation more effectively alters alcohol relapse-related behaviors, and whether these effects differ by sex.

Consistent with our original hypothesis, chemogenetic manipulation of the IL→NAcSh projection did not alter alcohol self-administration in females. These data might suggest that females rely less heavily on a single corticostriatal projection and instead recruit a broader network of reward- and affect-related circuits to regulate drinking. Recent circuit mapping studies demonstrate that binge-like alcohol exposure recruits more widespread nucleus accumbens inputs and broader patterns of neuronal activation in females than in males (Chan et al., 2025). Similarly, previous electrophysiological work from our laboratory demonstrated that CIA preferentially alters LFPs recorded in the central nucleus of the amygdala (CeA) in females, whereas corticostriatal circuits are more impacted by CIA in males (Hewitt et al., 2025). Although only one corticostriatal projection is examined in the present study, these converging findings support the broader hypothesis that alcohol dependence recruits distinct neural circuit strategies across the sexes, with females perhaps relying more heavily on distributed corticolimbic networks than males.

Disturbing activity within the IL→NAcSh pathway altered elevated plus maze (EPM) behavior in both sexes. Across exposure conditions, activation reduced open-arm time and increased closed-arm time in males, whereas both activation and inhibition reduced open-arm time in females. These changes in arm occupancy are consistent with greater open-arm avoidance, although reductions in distance traveled in CIA-exposed males and in females across exposure conditions complicate an anxiety-specific interpretation. The pattern of EPM effects diverged from the pattern of effects on alcohol self-administration. Inhibition increased alcohol intake during acute withdrawal in CIA-exposed males without a detectable change in open-arm time. In females, both manipulations reduced open-arm time without detectably altering alcohol intake. This dissociation suggests that the contributions of NAcSh-projecting IL neurons to anxiety-related behavior and withdrawal-associated drinking may be partly separable. The EPM data are just one snapshot of affective behavior, but highlight the complex interaction between negative emotional states and motivation. Addiction models propose that relief of withdrawal-related negative affect can sustain alcohol use (Koob, 2015), so future work should test whether manipulation of IL→NAcSh neurons changes the extent to which alcohol alleviates withdrawal-related negative affect to help clarify the role of this pathway in negative reinforcement mechanisms of AUD.

While the present study provides evidence that NAcSh-projecting IL neurons contribute to alcohol drinking differently in males and females, there are several methodological limitations to consider. The observed differences in IL c-Fos expression following DCZ administration were consistent with our intended chemogenetic effects, which provided strong support for our approach. However, c-Fos is an indirect marker of activity-dependent signaling and does not establish the magnitude or duration of changes in neuronal firing or their consequences for downstream circuits (Kovacs, 2008). Future studies combining chemogenetic manipulation with electrophysiological recordings or multisite fiber photometry could characterize how manipulating IL→NAcSh projections alters local activity and recruitment of interconnected regions during alcohol seeking.

Further, behavioral measures of anxiety should be interpreted cautiously. The absence of a CIA effect on open and closed arm time in the EPM may reflect methodological rather than biological factors, as mCherry controls were collected in a separate cohort that did not complete alcohol self-administration and therefore had somewhat different experiences across the approximately four-month experimental timeline. Secondly, although projection-specific DREADDs establish that the IL→NAcSh pathway contributes causally to behavior, they do not reveal the cellular or synaptic mechanisms responsible for those behavioral effects. Consequently, the present findings cannot determine whether the male-specific effects on alcohol drinking stem from altered glutamatergic transmission or inhibitory interneuron recruitment. Moreover, chemogenetic excitation produces relatively slow, sustained changes in neuronal excitability that differ substantially from the temporally precise firing patterns observed during endogenous circuit communication (Vlasov et al., 2018). Consequently, increasing tonic activity within the IL→NAcSh neurons may not reproduce the physiologically relevant patterns of activity required to suppress alcohol seeking, even if general neuronal excitability is successful. Future studies integrating chemogenetics with *ex vivo* electrophysiology, fiber photometry, calcium imaging, and molecular approaches will therefore be essential for identifying the cellular adaptations that underlie sex-specific corticostriatal dysfunction following chronic alcohol exposure.

Together, these findings suggest that intact IL→NAcSh activity supports behavioral control over drinking in alcohol-dependent males. Future studies should further investigate this male-specific effect by examining how IL→NAcSh signaling interacts with distributed networks involved in alcohol use and withdrawal; simultaneous recordings across multiple interconnected regions could clarify whether the behavioral consequences of IL→NAcSh manipulation reflect changes in coordinated activity across these broader networks. Identifying pathways that regulate alcohol drinking in dependent females is equally important. Our previous work implicates corticolimbic pathways in dependence-related phenotypes in females (Hewitt et al., 2025), highlighting the need to test their causal contributions to alcohol drinking. This work has important translational implications in that neuromodulation-based therapies continue to emerge as promising treatments for AUD and other substance use disorders (Bach et al., 2023; Davidson et al., 2022). The present findings raise the possibility that targeting the same circuit may have different therapeutic effects in males and females, and that understanding sex differences in circuit recruitment within distributed networks may help guide the development of more precise and biologically informed treatments for AUD.

